# High-quality genome assembly and functional annotation of Hiroshima-derived Pacific oyster *Magallana gigas*

**DOI:** 10.1101/2025.11.20.688662

**Authors:** Misako Imai, Miyuki Kanda, Moriya Mino, Kazuhiko Koike, Keisuke Okuhara, Hidemasa Bono

**Author notes:** Corresponding author: Hidemasa Bono.

## Abstract

The Pacific oyster (*Magallana gigas*) is a widely distributed marine bivalve of major importance in aquaculture and is abundant along the Asian coasts. The Hiroshima Prefecture is the principal region for oyster production in Japan. The currently available reference genomes are derived from oysters of UK origin and do not fully capture the genetic characteristics of the Hiroshima populations. Here, we present a high-quality genome assembly and functional annotation of *M. gigas* from Hiroshima, Japan. The genome was assembled using PacBio HiFi long reads. After assembly, the contaminant contigs were removed, yielding a final assembly with a total length of 707 Mb, contig N50 of 7.9 Mb, and BUSCO completeness of 99.9 %. Of the 44,639 predicted protein-coding genes identified using GALBA, 43,806 were successfully annotated using a functional annotation workflow (Fanflow). Overall, the high-quality assembly and annotation provide a valuable genomic resource for exploring the genetic diversity and environmental adaptations of Hiroshima oyster populations. The data will facilitate future studies in breeding, functional genomics, and genome editing.

## Introduction

The Pacific oyster (*Magallana gigas*) is an ecologically and economically important marine bivalve that is widely cultivated along the Asian coast. Recent studies and comprehensive reviews have shown that shellfish aquaculture plays a substantial biogeochemical role in coastal ecosystems; the Pacific oyster is a dominant filter feeder that contributes to water purification, nutrient cycling, and habitat stability^1,2^. Recent molecular studies have shown that Pacific oysters can rapidly adjust their immune and stress response pathways by activating pattern-recognition receptors, producing antimicrobial peptides, and upregulating heat-shock protein expression^3,4^. These responses help this species in tolerating marine pathogens and fluctuating environmental conditions encountered in aquaculture settings.

In Japan, the Hiroshima Prefecture is one of the major aquaculture centers for *M. gigas*, with well-established farming traditions and locally adapted breeding populations. Recently, Liu et al. presented the genome assembly of the closely related *Crassostrea* (*Magallana*) *sikamea* from Kumamoto, Japan, providing a valuable resource for comparative genomic studies on environmental adaptations within this genus^5^. Moreover, several reference genomes have been reported for *M. gigas*^6,7^, including assemblies derived from populations originating in the UK. However, these assemblies do not fully capture the genetic diversity and adaptive traits of Japanese oysters that have evolved under distinct environmental and aquaculture conditions. Despite their importance, no high-quality genome assemblies of Hiroshima oysters have been reported.

In the present study, we aimed to generate a high-quality genome assembly for a Hiroshima-derived Pacific Oyster using PacBio HiFi long-read sequencing. To establish a comprehensive genomic resource for this regionally specific population, we conducted gene structural and functional annotation. The dataset and annotation presented here provide an essential genomic foundation for future investigations into population structure, local adaptation, and immune-related gene evolution in Hiroshima oysters. They may also support breeding strategy that enhance the ability of oysters to adapt to rapidly changing aquaculture environments.

## Methods

### Ethics statement

Pacific oyster (*M.gigas*) used in this study was obtained from commercial aquaculture company located in Akitsu, Higashi-Hiroshima, Hiroshima, Japan. As oysters (*M.gigas*) are invertebrates and not subject to animal experimentation regulations in Japan, no specific ethical approval was required for this study.

### Sample collection and DNA sequencing

Seven-month-old oysters were collected from aquaculture farms of Mitsu Bay in Akitsu, Higashi-Hiroshima, Hiroshima Prefecture, Japan (34.31°N, 132.80°E) (Figure 1a). Genomic DNA was extracted from adductor muscle tissue of a single individual using the NucleoBond HMW DNA kit (Macherey-Nagel, Düren, Germany), according to the manufacturer’s instructions. The quality and quantity of the extracted genomic DNA were assessed using a Qubit 4 fluorometer (Thermo Fisher Scientific, Waltham, MA, USA) and Femto Pulse system (Agilent Technologies, Santa Clara, CA, USA). DNA was sheared into 10–30 kb fragments using a g-TUBE device (Covaris Inc., Woburn, MA, USA). A single-molecule real-time (SMRT)bell library was constructed for HiFi sequencing using the SMRTbell Prep Kit version 3.0 (Pacific Biosciences, Menlo Park, CA, USA) following the manufacturer’s protocol. The library was sequenced using the PacBio Sequel IIe platform (SMRT Cell 8M; Sequel II Binding Kit version 3.2). An overview of the workflow of genome analysis is shown in Figure 1b. K-mer analysis (k = 31) was performed using GenomeScope version 2.0^8^.

**Figure 1.**
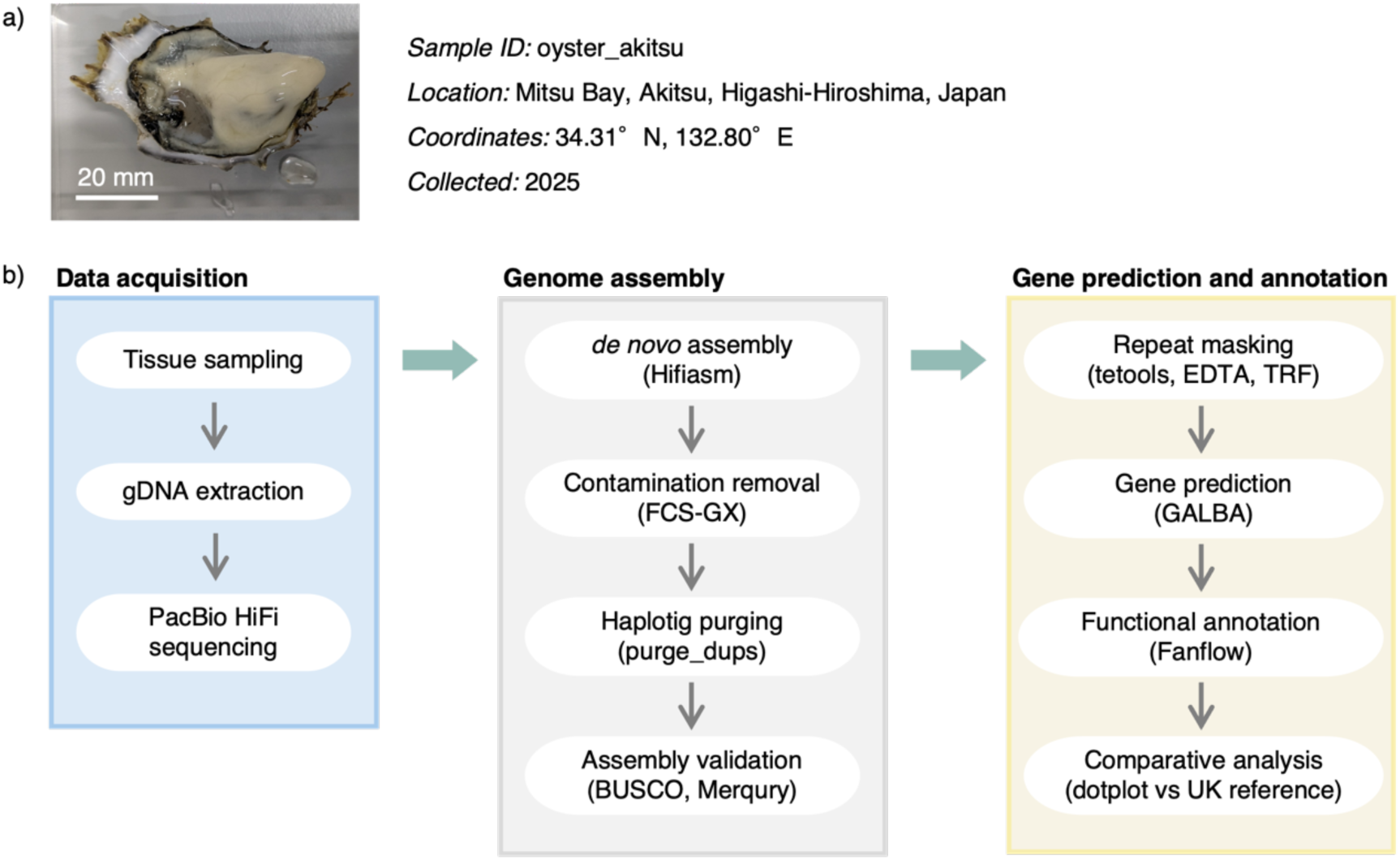
Overview of the sample collection and genome analysis workflow for Hiroshima-derived *Magallana gigas*. (**a**) Photograph of the seven-month-old oyster collected from Mitsu Bay, Akitsu, Higashi-Hiroshima, Japan (34.31°N, 132.80°E). (**b**) The genome analysis workflow was divided into three stages: (1) data acquisition, including DNA extraction and PacBio HiFi sequencing; (2) genome assembly, including contamination removal, haplotig purging, and quality assessment; and (3) gene prediction and functional annotation.

### Genome assembly

The Hifiasm (version 0.22.0)^9–11^ “--primary” option was used for *de novo* assembly, generating primary (p_ctg) and alternate (a_ctg) contigs. FCS-Gx (version 0.5.4)^12^ was used to screen and remove for potential contaminants from the primary assembly. The haplotigs were purged using purge_dups^13^ (default parameters). Basic assembly statistics were obtained using SeqKit (version 2.10.0)^14^, and assembly completeness was assessed using BUSCO (version 5.7.1; mollusca_odb10; genome mode). Base-level consensus accuracy was estimated using Merqury (version 1.3)^15^. To verify assembly consistency, PacBio HiFi reads were realigned to the assembly using minimap2 (version 2.28)^16^ with the preset -x map-hifi, and mapping statistics were summarized using SAMtools flagstat (version 1.19.2)^17^.

### Repeat masking

Repetitive elements were identified using EDTA version 2.1.0^18^ and RepeatModeler^19^, which are included in Dfam TEtools version 1.88.5^20^, and the repeat-masking strategy described in the GALBA study^21^ was followed. Consensus sequences obtained using both methods were merged. Redundant sequences were removed using CD-HIT-EST version 4.8.1^22,23^, at 90 % sequence identity, to generate the initial non-redundant repeat library. The genome was first soft-masked with RepeatMasker (included in Dfam TEtools version 1.88.5) using this library. Subsequently, tandem repeats were detected using TRF^24^ and masked onto the soft-masked genome. Subsequently, the TRF-derived tandem repeat library was combined with the initial repeat library, and redundancy was again removed using CD-HIT-EST at 90 % identity to produce the final composite repeat database.

### Gene and functional annotations

Protein-coding genes were predicted using GALBA version 1.0.11.2^21^, which applies an ab initio approach supported by evidence on protein homology. The completeness of the predicted gene models was assessed using BUSCO version 5.7.1^25,26^ (mollusca_odb10; protein mode). Functional annotation of predicted protein-coding genes was performed using Fanflow^27^. Amino acid sequences were compared individually with the annotated protein datasets using ggsearch36 (included in fasta-36.3.8i^28^). The protein domains were identified using HMMSCAN (HMMER^29^ version 3.3.2) with Pfam version 37.3 as the domain database.

### Comparative genome alignment

The UK-derived reference genome (xbMagGiga1.1; GCA_963853765.1) was downloaded from the NCBI Genome database. Basic assembly statistics were recalculated using SeqKit and BUSCO to ensure consistency of the Hiroshima-derived genome sequence. For comparative analysis, sequences shorter than 1 Mb were excluded from both assemblies before alignment. Pairwise genome alignment was performed using minimap2 (version 2.28) with the -x asm5 option. The resulting PAF alignment file was visualized as a dot plot using dotPlotly^30^ (default settings) to assess large-scale structural correspondence between the two assemblies.

## Results and discussion

### Read quality and k-mer profiling

Approximately 2.3 million HiFi reads were generated, totaling 38.6 Gbp, with an average read length of 16.7 kb. K-mer profiling was performed using GenomeScope version 2 (k = 31) with the PacBio HiFi reads. The estimated haploid genome size was approximately 471 Mb with a heterozygosity rate of 3.36 % (Figure 2). The resulting HiFi reads were used for genome assembly.

**Figure 2.**
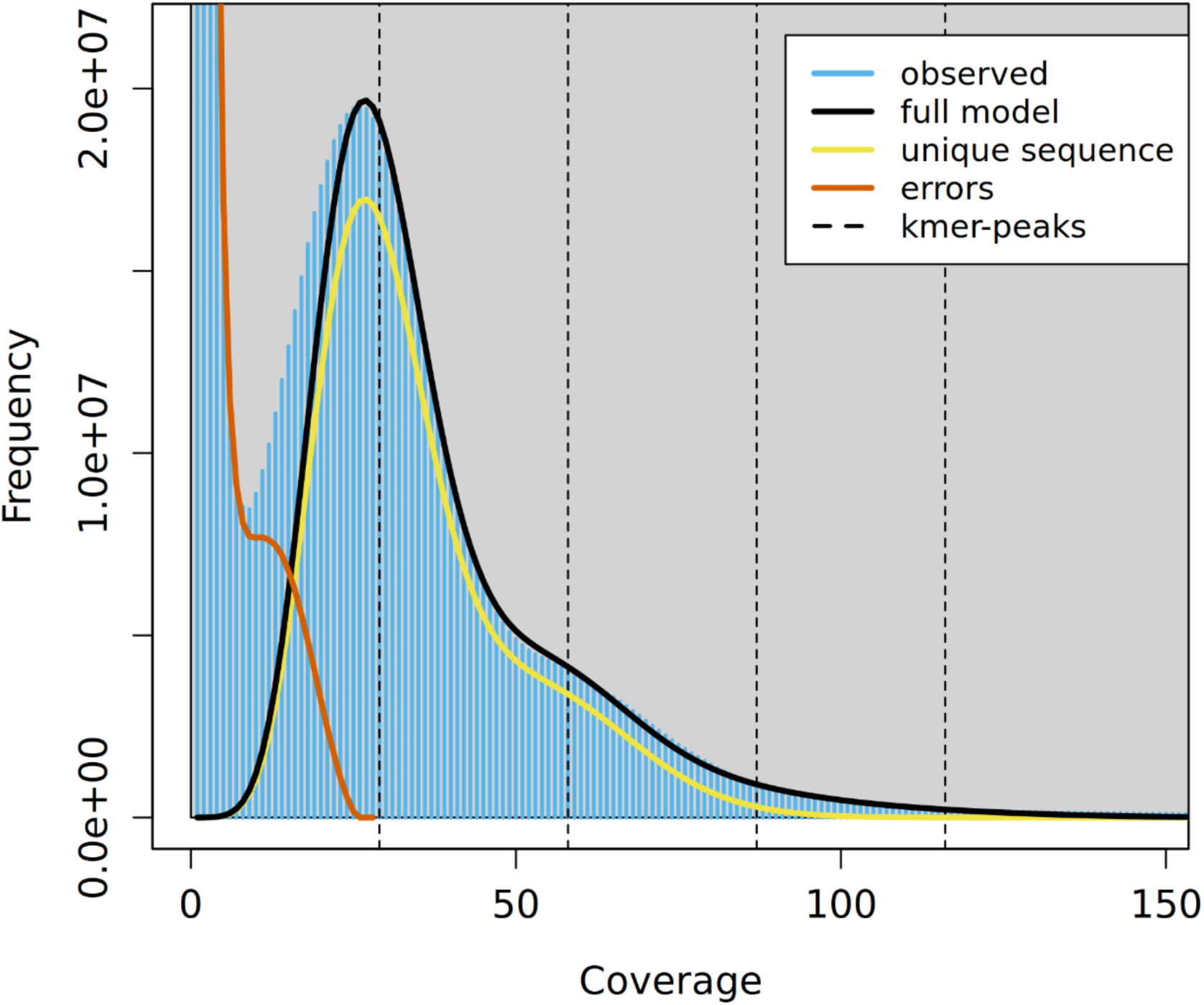
K-mer distribution and genome profiling using GenomeScope version 2. K-mer frequency histogram (k = 31) generated from PacBio HiFi reads, modeled using GenomeScope version 2. The estimated haploid genome size was approximately 471 Mb, with a heterozygosity rate of 3.36 %, indicating that it was a diploid genome with moderate heterozygosity. The first (lower-coverage) peak represents heterozygous k-mers, whereas the second (higher-coverage) peak corresponds to homozygous k-mers.

### Genome assembly

De novo assembly was performed using Hfiasm (version 0.22.0), generating primary (p_ctg) contigs. The draft assembly was screened for potential contamination using FCS-GX (GenomeClean version 0.5.4); one contig comprised a prokaryotic viral sequence similar to that of a *Vibrio* phage (*Vibrio* phage 252E42.2), and thus, the contig was removed. Because oysters often host microbiota, among which *Vibrio* species are predominant^2,3,31^, the detection of *Vibrio* phage-related sequences is likely due to environmental contamination. No additional contaminants were detected after removal. The final primary assembly spanned 707 Mb with contig N50 of 7.9Mb (Table 1). Assembly completeness was assessed using BUSCO (version 5.7.1; mollusca_odb10; genome mode), which showed 99.9 % completeness (single-copy, 94.3 %; duplicated, 5.6 %; fragmented, 0.2 %; missing, 0.1 %). Base-level consensus accuracy was estimated using Merqury (version 1.3)^15^, yielding a quality value (QV) of 60.34 for the primary contigs (p_ctg), corresponding to an error rate of less than 10⁻⁶. The Hiroshima-derived assembly comprises contigs, and therefore, it is less continuous than the chromosome-scale UK-derived reference genome; the high level of BUSCO completeness (99.9 %) suggested a near-complete gene representation in the Hiroshima-derived assembly comparable to or exceeding that of the existing reference genome.

**Table 1.**
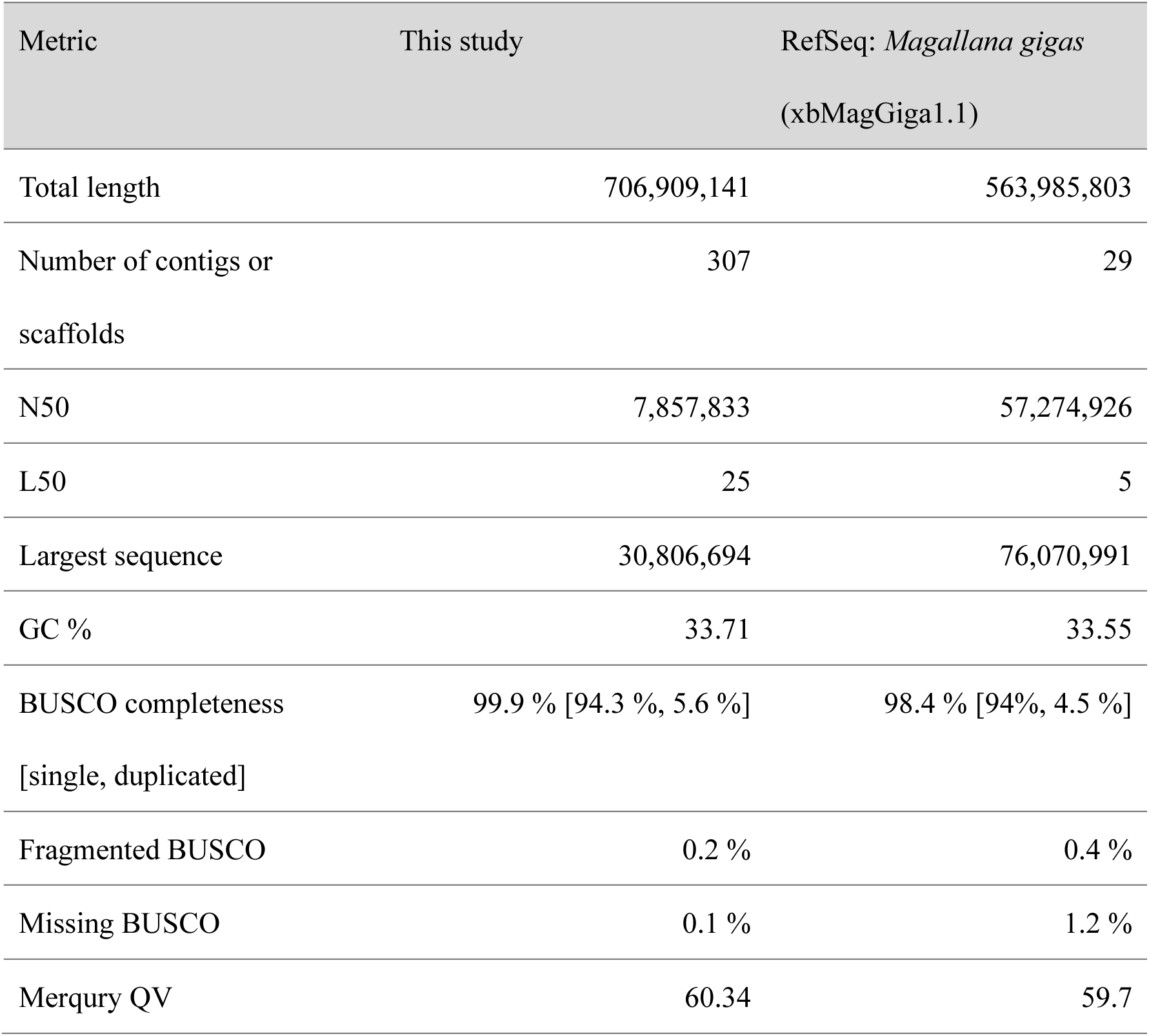
Comparison between genome assembly statistics of the Hiroshima-derived *Magallana gigas* genome and the UK-derived reference (xbMagGiga1.1).

The PacBio HiFi reads were mapped to the final assembly using minimap2 (-x map-hifi), resulting in an average alignment rate of 99.8 %, confirming that approximately all the input reads were properly represented in the assembly.

### Annotation of Hiroshima-derived *M.gigas* genome

A total of 53.2 % of the genome was identified as repetitive or masked, and this soft-masked assembly was subsequently used as the input for gene prediction. The gene prediction was performed in GALBA ab initio mode (--AUGUSTUS_ab_initio). A custom protein database for five closely related oyster species retrieved from NCBI was also incorporated, as shown in Table 2. As a result, 44,639 protein-coding genes was predicted (Table 3). Annotation-level completeness was evaluated by examining the predicted gene set using BUSCO in the protein mode; consequently, 98.6 % of the expected single-copy orthologs (single, 69.3 %; duplicated, 29.3 %; fragmented, 0.4 %; missing, 1 %) were recovered, indicating a high coverage of conserved genes.

**Table 2.** Reference protein datasets used for gene prediction using GALBA.

| Species | Scientific name | Database / Source | Accession |
| --- | --- | --- | --- |
| Pacific oyster (UK-derived reference) | *Magallana gigas* | NCBI Genome | GCF_963853765.1 |
| Eastern oyster | *Crassostrea virginica* | NCBI Genome | GCF_002022765.2 |
| Portuguese oyster | *Magallana angulata* | NCBI Genome | GCF_025612915.1 |
| Tropical rock oyster | *Saccostrea echinata* | NCBI Genome | GCF_033153115.1 |
| European flat oyster | *Ostrea edulis* | NCBI Genome | GCF_947568905.1 |

**Table 3.** Comparison between gene prediction statistics of the Hiroshima-derived genome and UK-derived *Magallana gigas* (xbMagGiga1.1) reference genome.

| Metric | This study | Ensembl metazoa release 62<br>(xbMagGiga1.1) |
| --- | --- | --- |
| Number of protein-coding<br>genes | 44,639 | 19,775 |
| Number of transcripts | 53,768 | 37,189 |
| BUSCO completeness<br>[single, duplicated] | 98.6 % [69.3 %, 29.3 %] | 96.5 % [54.1 %, 42.4 %] |
| Fragmented BUSCO | 0.4 % | 0.2 % |
| Missing BUSCO | 1 % | 3.3 % |

For functional annotation, we applied Fanflow using species-specific protein reference datasets and Pfam-based domain assignments (Table 4). Of the predicted genes, 43,806 (98.1%) were successfully annotated based on orthological data and protein domain evidence (Table 5). Taken together, these results confirm that the Hiroshima oyster genome assembly exhibits a high degree of completeness, base accuracy, and structural reliability, thereby providing a robust reference resource for downstream comparative and functional genomic analyses

**Table 4.** Protein and domain reference datasets used for functional annotation with Fanflow.

| Species / Dataset | Scientific name | Database / Source | Accession |
| --- | --- | --- | --- |
| Pacific oyster (UK-derived reference) | *Magallana gigas*<br>(xbMagGiga1.1) | Ensembl Metazoa<br>(release 61) | GCA_963853765.1 |
| Pacific oyster | *Magallana gigas*<br>(uk_roslin_v1) | Ensembl Metazoa<br>(release 61) | GCA_902806645.1 |
| Eastern oyster | *Crassostrea virginica* | Ensembl Metazoa<br>(release 61) | GCA_002022765.4 |
| Portuguese oyster | *Magallana angulata* | Ensembl Metazoa<br>(release 61) | GCA_025612915.2 |
| Worm | *Caenorhabditis elegans* | Ensembl (release 113) | GCA_000002985.3 |
| Mouse | *Mus musculus* | Ensembl (release 113) | GCA_000001635.9 |
| Human | *Homo sapiens* | Ensembl (release 113) | GCA_000001405.28 |
| UniprotKB/Swiss-Prot (protein reference) | - | UniProt (release 2025_01) | - |
| Pfam (protein domain reference) | - | EMBL-EBI / InterPro | Pfam release 37.3 |

**Table 5.** Summary of protein-coding gene prediction and annotation for the Hiroshima-derived *Magallana gigas* genome.

| Annotation category | Annotation level | No. of genes |
| --- | --- | --- |
| Protein homologue from tophit | *Magallana gigas* (xbMagGiga1.1) | 36,456 |
|  | *Magallana gigas* (uk_roslinv1) | 39,034 |
|  | *Magallana angulata* | 37,560 |
|  | *Crassostrea virginica* | 35,995 |
|  | *Caenorhabditis elegans* | 19,052 |
|  | *Drosophila melanogaster* | 17,934 |
|  | *Mus musculus* | 24,746 |
|  | *Homo sapiens* | 25,878 |
|  | UniProtKB/Swiss-Prot | 27,686 |
|  | At least one of the above | 43,714 |
| No protein homolog | Protein domain | 92 |
|  | At least one of the above | 43,806 |
| Hypothetical protein | — | 833 |
| Total | — | 44,639 |

### Comparative genome analysis

To assess large-scale structural consistency, pairwise alignment between Hiroshima-derived contigs and the UK-derived chromosome-scale *M. gigas* reference genome was performed using minimap2 (-x asm5) and visualized using dotPlotly. The dot plot revealed an overall consistency in sequence alignment, with several locally redundant regions that may correspond to segmental duplications or copy number variations (CNVs) (Figure 3).

**Figure 3.**
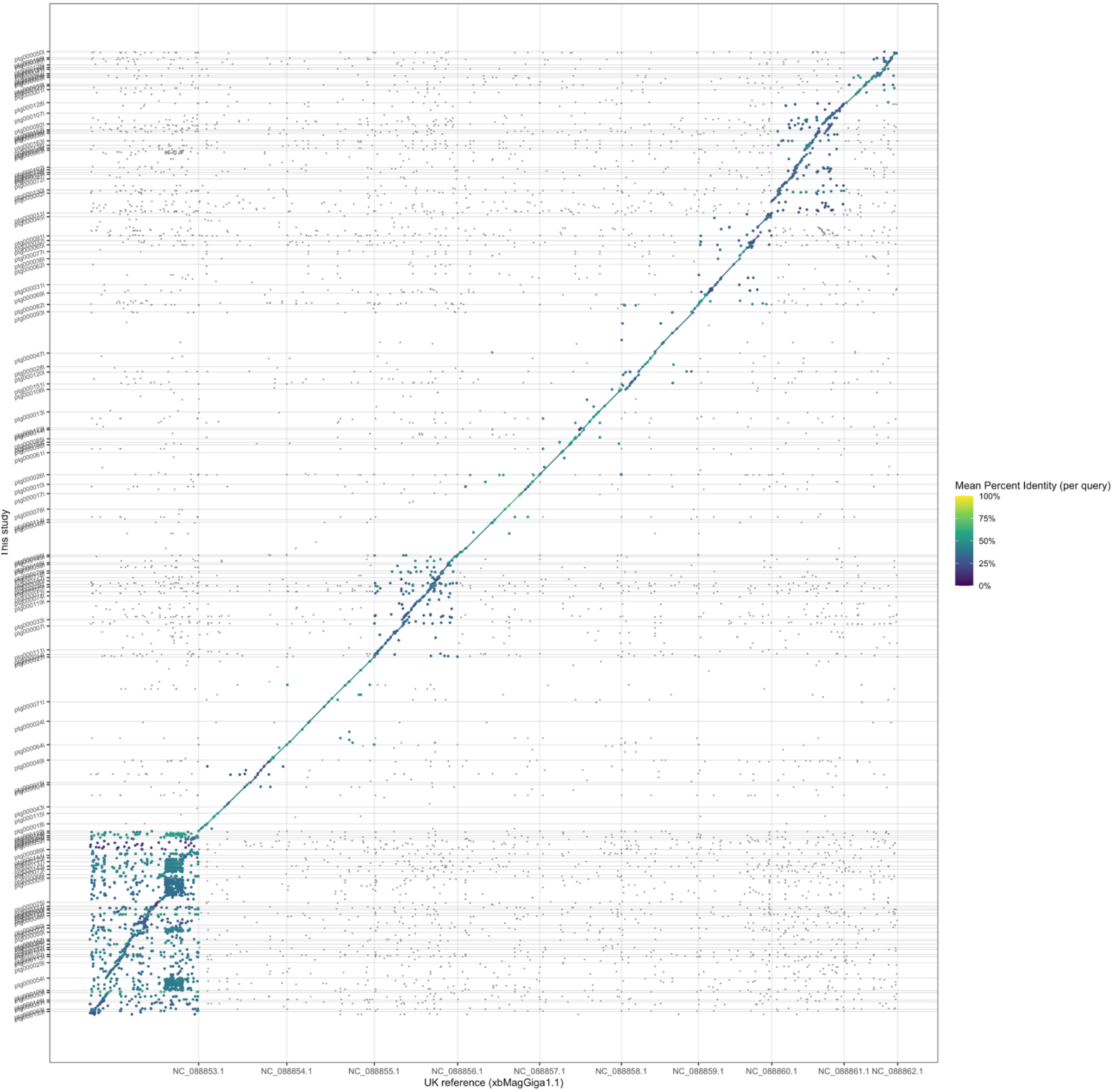
Comparative alignment of the Hiroshima-derived and UK reference (xbMagGiga1.1) genomes. Dot plot generated using minimap2 (-x asm5) and visualized with dotPlotly, comparing the Hiroshima-derived contigs with the chromosome-scale UK reference genome. The dot plot shows largely concordant sequence alignments between the Hiroshima-derived and UK reference genomes. The dot plot also shows locally redundant regions.

## Data availability

All datasets generated and analysed in this study are publicly available. Raw PacBio HiFi sequencing reads and assembled genome sequences have been deposited in the DNA Data Bank of Japan (DDBJ) under the BioProject accession number PRJDB38101. Additional resources are provided on figshare, including: (i) the repeat library, repetitive sequence statistics, and masked genome^32^; (ii) predicted gene models in GFF3, protein, and transcript FASTA formats^33^; and (iii) functional annotation resources generated using Fanflow^34^.

## Code availability

The modified dotPlotly code used in this study is available on figshare^35^. The visualization script used for constructing comparative genome dot plots was adapted from dotPlotly (MIT License), with minor modifications to improve visual clarity. All other analyses were performed using publicly available software as described in the Methods section.

## Author contributions

H.B. and K.O. coordinated and designed this study, and K.K. and M.M. collected the Pacific oyster samples. M.K. conducted the sampling and sequencing. M.I. and H.B. performed the bioinformatic analyses and prepared figures and tables. M.I. wrote the first draft of the manuscript. All authors reviewed and approved the final manuscript.

## Acknowledgements

Computational analyses were performed using computing resources provided by the Bio-DX Organization and internal servers at PtBio Inc. We thank all members of the laboratory at Hiroshima University and the R&D division of PtBio Inc. for their collaboration and valuable support.

